# Latent transforming growth factor binding protein-2 (LTBP2), an IPF biomarker of clinical decline, promotes TGF-beta signaling and lung fibrosis in mice

**DOI:** 10.1101/2025.05.06.652563

**Authors:** Nicholas K Bodmer, Malay Choudhury, Haider Mirza, Suramamayi Pradhan, Yongjun Yin, Robert P Mecham, Steven L Brody, David M Ornitz, Jeffrey R Koenitzer

## Abstract

The identification of clinically predictive serum biomarkers for pulmonary fibrosis is a significant challenge and important goal. Multiple recent proteomic biomarker studies have identified latent transforming growth factor binding protein-2 (LTBP2) as a circulating factor associated with disease progression in fibrotic lung diseases in humans (including IPF), but its role in the development of fibrosis is incompletely defined. LTBP2 competes with the large latent transforming growth factor-beta (TGFβ) complex (LLC) for binding to the N-terminus of fibrillin and is thought to promote the release of active TGFβ. We hypothesized that LTBP2 deficiency would promote LLC sequestration in matrix and reduce TGFβ signaling. We recently reported an LTBP2 knockout (*Ltbp2^-/-^*) mouse with no baseline lung abnormalities. Here we show that *Ltbp2^-/-^* mice exposed to either bleomycin or silica have a significant reduction in fibrosis compared to wild type controls. Consistent with reduced fibrosis, after bleomycin *Ltbp2^-/-^* mouse lungs have reduced TGFβ signaling and isolated fibroblasts from *Ltbp2^-/-^*mice exhibit impaired migration in an in vitro wound closure assay. Transcriptomic analysis of bleomycin-treated control and *Ltbp2^-/-^* mouse lung tissue identified multiple LTBP2-regulated genes, including the lncRNA antisense of IGFR2 non-coding RNA (*Airn*) which has reported antifibrotic effects. Interestingly, we also observed that *Ltbp2^-/-^* mice had impaired epithelial repair after bleomycin treatment, a phenotype that also occurred in a naphthalene model of club cell injury. These findings provide evidence that LTBP2 is profibrotic and facilitates TGFβ signaling but is also required for normal airway epithelial repair.

## Introduction

Pulmonary fibrosis is the pathologic endpoint of multiple interstitial lung diseases whose unifying feature is an aberrant response to lung injury resulting in excessive collagen deposition^1^. At homeostasis, the signaling of growth factors involved in the fibrotic response is tightly regulated through their binding to extracellular matrix (ECM) components^2–4^. In pulmonary fibrosis, the composition of ECM is fundamentally altered with fibrotic lung regions marked by increased expression of various proteoglycans, fibrillar collagens, fibulins and fibrillins, and latent transforming growth factor binding proteins (LTBPs)^5^. Identifying ECM proteins that facilitate signaling of pro-fibrotic growth factors may enable disruption of these interactions as a therapeutic approach.

Latent transforming growth factor binding protein-2 (LTBP2) is a secreted microfibril-interacting protein synthesized by fibroblasts^6^. Unlike other members of the LTBP family, LTBP2 does not bind transforming growth factor-beta (TGFβ) but may indirectly influence its signaling in a variety of ways, including competition for the large latent complex (LLC) binding site near the N-terminus of fibrillin-1^7,8^. LTBP2 expression is increased in fibrotic lung disease and its overexpression in cultured fibroblasts is sufficient to induce myofibroblast differentiation^9,10^, but a gap in knowledge persists regarding *in vivo* mechanisms of pro-fibrotic activity.

Given the variable clinical course of pulmonary fibrosis, a focus of clinical research is identification of biomarkers to recognize patients likely to experience rapidly progressive disease. Multiple recent independent studies identified LTBP2 as a serum biomarker for pulmonary fibrosis associated with poor clinical outcomes, including a large unbiased screen in which LTBP2 was the strongest negative predictor of transplant-free survival of all measured proteins, with a hazard ratio of 2.43^11–13^. Understanding the function of LTBP2 in the fibrotic milieu would increase its clinical value as a serum biomarker, as it may allow the selective deployment of targeted therapies to high-risk patient groups.

We previously generated and characterized LTBP2 null (*Ltbp2*^-/-^) mice, which provide a specific tool for identifying LTBP2-regulated pathways^14^. Here, we use multiple *in vivo* lung injury models to delineate LTBP2 function in pulmonary fibrosis. We show that *Ltbp2*^-/-^ mice have reduced injury-induced fibrosis due to impaired TGFβ signaling. Using bulk RNA sequencing (RNASeq) and airway-focused immunostaining, we identify an additional role for LTBP2 in facilitating repair of the airway epithelium following lung injury.

## Methods

### Human lung tissues

Studies of anonymized tissues from lungs of deceased individuals donated for research and those explanted during lung transplantation were approved by the Institutional Review Board at Washington University. Non-disease donors were those unsuitable for lung transplantation. Human lung fibroblasts isolated from non-diseased lungs donated for research and from lungs explanted from patients with idiopathic pulmonary fibrosis were obtained as previously described^15^.

### Mouse models of lung injury

Studies were approved by the Institutional Animal Care and Use Committee at Washington University. Male and female C57Bl/6J wild-type mice (Jackson Labs), 8-12 weeks of age, or *Ltbp2^-/-^* mice (C57Bl/6 background), 20-25 grams, were used for all models. Bleomycin was administered intranasally at 1.5U/kg, silica oropharyngeally (10mg), and naphthalene intraperitoneally (250 mg/kg) as previously described^16,17^. At experimental endpoints mice were euthanized with ketamine and xylazine and lungs were snap frozen or pressure fixed via 10% phosphate-buffered formalin. After overnight fixation samples were dehydrated, submitted for paraffin embedding, and cut in to 5-μm sections. Samples were stained with H&E or Masson’s trichrome as described^16^.

### Immunohistochemistry and immunofluorescence

After deparaffinization with xylene or citric acid, sections were rehydrated in sequential ethanol baths. Immunochemistry or immunofluorescence were then performed using standard techniques. Further details are included in the Materials and Methods online supplement.

### Primary fibroblast culture and scratch assay

Isolated lungs were treated with dispase and collagenase, minced on a petri dish, then gently shaken at 37 °C for 1 h. Dissociated cells were passed through sterile cell strainers, and centrifuged. The pellet was resuspended and incubated on a 75 mm petri dish. Cells were re-plated in 48 well plates and once confluent, scratched with a sterile P200 pipet tip. The defect width was measured in 3 locations. In some experiments the TGFβ receptor inhibitor SB431542 (Tocris) was included.

### Fibrosis quantification

Trichrome-stained lung sections were scanned on a Zeiss Axio Scan.Z1 Slide Scanner at 20× magnification. Qupath software (version 0.4.3) was used to extract TIFF files from whole-slide images. For bleomycin-induced fibrosis, sections were divided into 2-3 similar size regions, to include the entire lobe, then scored by two blind readers using the modified Ashcroft score criteria^19^. For silica-induced fibrosis, Qupath was used to quantify percentage of lung covered by fibrotic masses after exclusion of bronchi.

### RNA Sequencing

Lung tissue was stored at −80 °C until processing. mRNA isolation was performed with the Ambion PureLink RNA Mini Kit (Thermo Fisher, 12183018, Lot 2379554), and samples submitted for library preparation, sequencing, and analysis through the Washington University Genome Technology Access Center (GTAC), as previously described^20^. Details are available in the Materials and Methods online supplement.

### Single cell RNASeq analysis

Datasets GSE132771 (fibrosis) and GSE173896 (COPD) were accessed via the Gene Expression Omnibus (GEO) and analyzed using standard *Seurat* pipelines. Details are available in the Materials and Methods online supplement.

### Statistical Analysis

Significant differences in mean values were calculated using paired Student’s *t* tests or one-way ANOVA. Mortality was analyzed using log-rank (Mantel-Cox) test and two-way ANOVA. A *P* value of less than 0.05 was considered significant.

## Results

### LTBP2 is a marker of pathologic fibroblasts and tissue remodeling in parenchymal and airway fibrosis

Previous studies have reported increased LTBP2 protein in lung tissue from patients with IPF and COPD, in sorted myofibroblasts from mice after bleomycin, and in induced myofibroblasts from human lungs^9^. As recent single cell RNASeq analyses have improved resolution of mesenchymal cell subpopulations, we used two publicly available single cell RNA sequencing datasets to evaluate LTBP2 mRNA expression in pulmonary fibrosis and COPD^21,22^. Collagen triple helix repeat containing 1 (*Cthrc1*)-positive pathologic fibroblasts expand in both pathologies (though to a greater extent in pulmonary fibrosis) and are enriched for LTBP2 expression compared to other fibroblast populations (Fig 1A-D, S1A,B), though LTBP2 expression per cell is generally unchanged with disease in these datasets (not shown). To confirm that LTBP2 expression is increased in remodeled regions of diseased lung, we used immunofluorescence microscopy in control donor lungs, end-stage explants of IPF and COPD lungs, and donor lungs with severe asthma. In representative samples of at least three different lungs, LTBP2 protein expression was minimal in control lungs (Fig 1E) but increased in each disease, marking parenchymal fibrosis in IPF and remodeled airways in COPD and asthma (Fig 1F-H). To further confirm the cellular source of LTBP2 in fibrosis we performed immunoblot analysis for LTBP2 in primary fibroblasts derived from IPF versus control lungs and observed increased LTBP2 expression in IPF-derived cells (Fig 1I, J).

**Figure 1.**
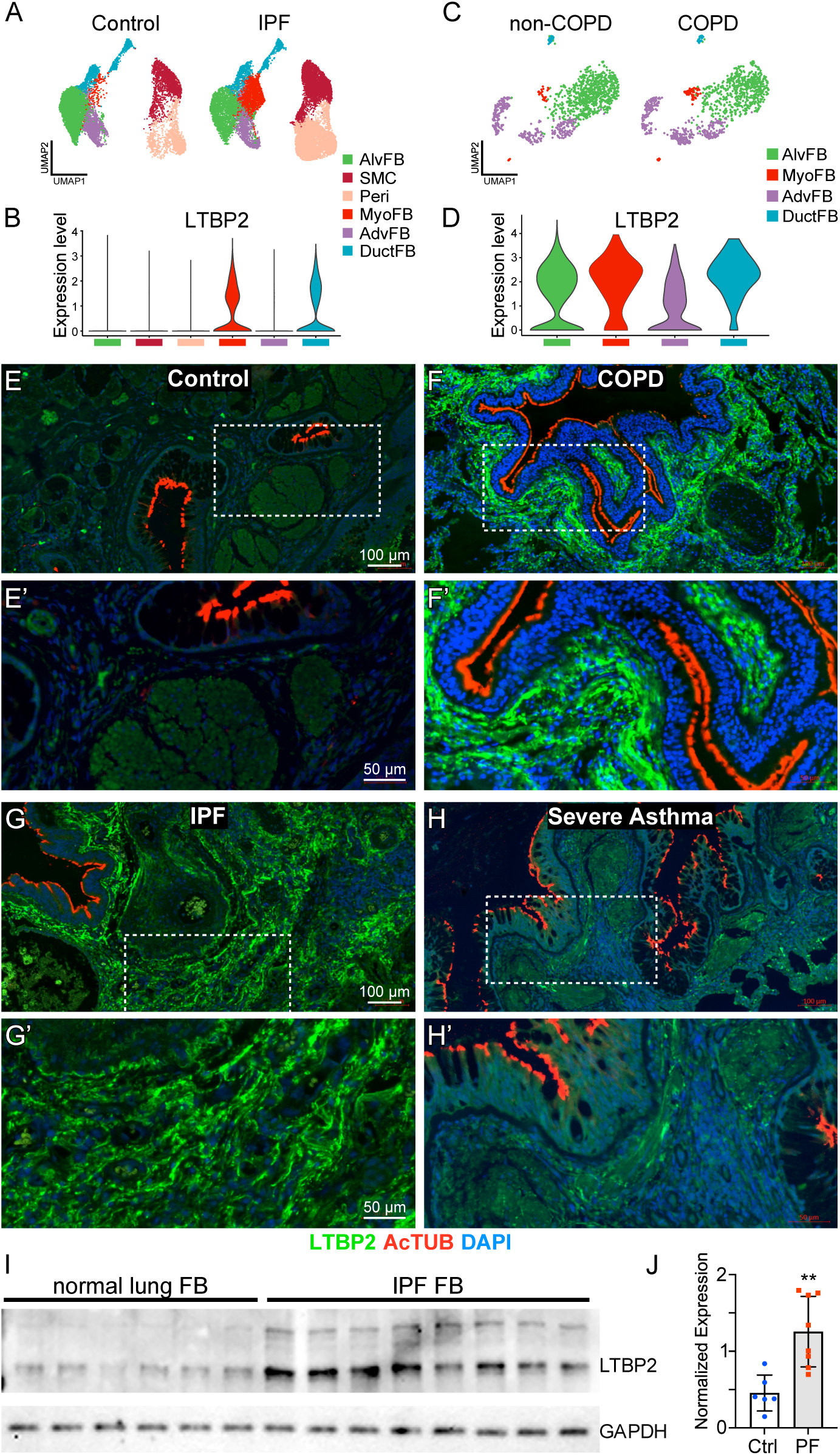
LTBP2 is expressed in pathologic fibroblasts and increased in remodeled lung. (A) UMAP with scRNASeq clusters of mesenchymal subpopulations from fibrotic and control lungs. (B) Violin plot with expression levels of *LTBP2* within mesenchymal subpopulations from 1A. (C) UMAP with scRNASeq clusters of fibroblast subpopulations from COPD and control lungs. (D) Violin plot of *LTBP2* expression levels within mesenchymal subpopulations from 1C. (E) Immunofluorescence of lung sections from human donor control lung with LTBP2 (green), acetylated tubulin (red), and DAPI (blue), with highlighted area magnified in E’. (F) Immunofluorescence of lung sections from end-stage COPD explanted lung with LTBP2 (green), acetylated tubulin (red), and DAPI (blue), with highlighted area magnified in F’. (G) Immunofluorescence of lung sections from end-stage IPF explanted lung with LTBP2 (green), acetylated tubulin (red), and DAPI (blue), with highlighted area magnified in G’. (H) Immunofluorescence of lung sections from severe asthma donor with LTBP2 (green), acetylated tubulin (red), and DAPI (blue), with highlighted area magnified in H’. (I) Western blot depicting LTBP2 and GAPDH bands for primary fibroblasts derived from control lungs or IPF explant lungs. n=6 non disease control; n=8 IPF unique explanted lungs. (J) Quantification of LTBP2 expression per group (normalized to GAPDH) from (I).

### LTBP2-null mice are protected from bleomycin- and silica-induced fibrosis

Two inhalational models, bleomycin and silica, were used to assess the role of LTBP2 in fibrosis, with tissue collection at multiple time points (Fig 2A). In male mice in the bleomycin model, LTBP2 deficiency conferred a survival advantage, while mortality in female bleomycin-treated mice was not significantly altered by bleomycin injury (Fig 2B). *Ltbp2*^-/-^ mice in the bleomycin model had reduced fibrosis burden at 28 d by histology (Fig 2C) as quantified by Ashcroft scoring (Fig 2D) and reduced collagen content by hydroxyproline assay (Fig 2E). Collagen visualization by picrosirius red stain under polarized light (Fig 2F) showed reduced collagen deposition in *Ltbp2*^-/-^ lungs as early as 14 d. While no early mortality was observed in either group in the silica model (Fig 2G), we again observed a significant reduction in the extent of fibrosis at 28 d in trichrome stained sections (Fig 2H,I).

**Figure 2.**
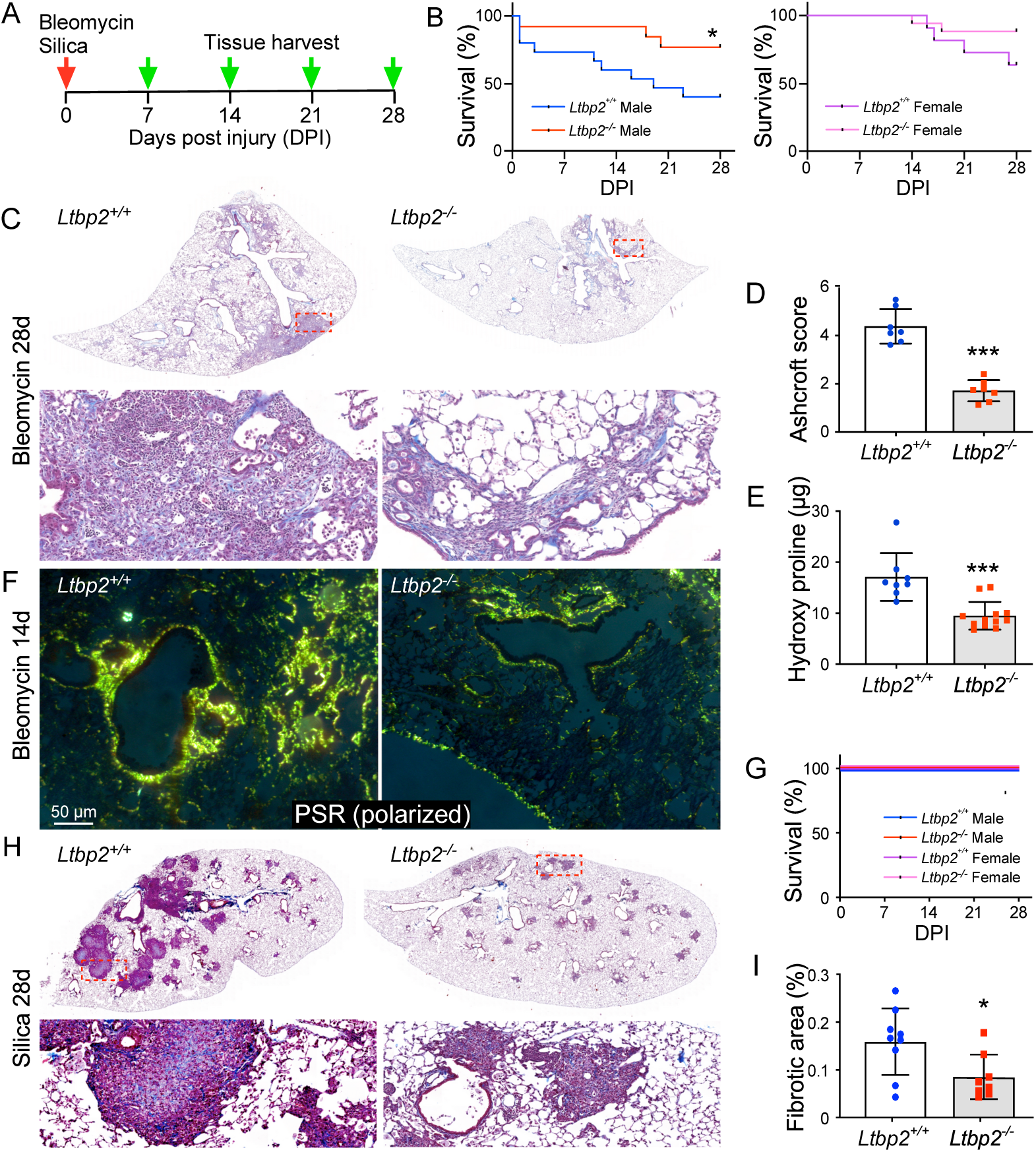
LTBP2 null mice are protected from lung fibrosis. (A) Scheme of silica (oropharyngeal, 10 mg) and bleomycin (intranasal, 1.6 U/kg) administration and tissue collection time course for mice. (B) Kaplan-Meier curves for male (L) and female (R) mice, *Ltbp2*^-/-^ versus WT. (C) Representative Trichrome-stained sections of bleomycin-treated lungs at 28d, *Ltbp2*^+/+^ versus *Ltbp2*^-/-^ mice. (D) Bar graph of blinded Ashcroft score averages per individual mouse (n=7 per group). (E) Bar graph of hydroxyproline levels measured in *Ltbp2*^-/-^ and WT lungs (n=7 per group). (F) Polarized microscopy of picrosirius red in bleomycin treated *Ltbp2*^-/-^ and WT lungs at 14 d. (G) Kaplan-Meier curve depicting survival of mice in oropharyngeal silica model. (H) Representative Trichrome-stained sections of silica-treated lungs at 28 d from *Ltbp2*^+/+^ versus *Ltbp2*^-/-^ mice at low power (top panels) vs. higher power (lower panels). (I) Bar graph depicting measured fibrotic area, average per section per mouse, n=9.

### TGFβ signaling and development of pathologic fibroblasts are abrogated in LTBP2-null mice

Similar to human lungs, minimal protein expression of LTBP2 was observed in naïve control conditions (Fig 3A). In bleomycin-induced fibrosis, we noted increased LTBP2 protein expression (with absent expression in *Ltbp2*^-/-^ mice) in lung regions also marked by expression of the pathologic fibroblast markers alpha-smooth muscle actin (⍺SMA) or CTHRC1 (Fig 3B,C). CTHRC1 expression was also reduced, even in remodeled lung regions in *Ltbp2*^-/-^ mice (Fig 3D). Quantitative RT-PCR (qRT-PCR) also showed a reduction in *Acta2* expression (Fig 3E), suggesting reductions in myofibroblast and overall fibroblast burden. Given that large latent TGFβ complex binding and sequestration by microfibrils may be subject to competition by LTBP2, we hypothesized that the observed reduction in fibrosis in *Ltbp2*^-/-^ mice would be associated with a reduction in TGFβ signaling. Immunohistochemistry for phospho-SMAD2 and SMAD3 were not altered in untreated lungs from *Ltbp2^-/-^* mice (Fig 3F), but both markers were decreased in *Ltbp2*^-/-^ mice at 28 days post-bleomycin injury (Fig 3G,H). The reduction in pSMAD2 as a fraction of total SMAD2 was confirmed by immunoblot analysis and was accompanied, as expected, by a reduction in collagen 1 protein (Fig 3I). Comparison was again made to the silica model, where we observed reductions in pSMAD2 and pSMAD3, especially within organized nodules (Fig 3J,K).

**Figure 3.**
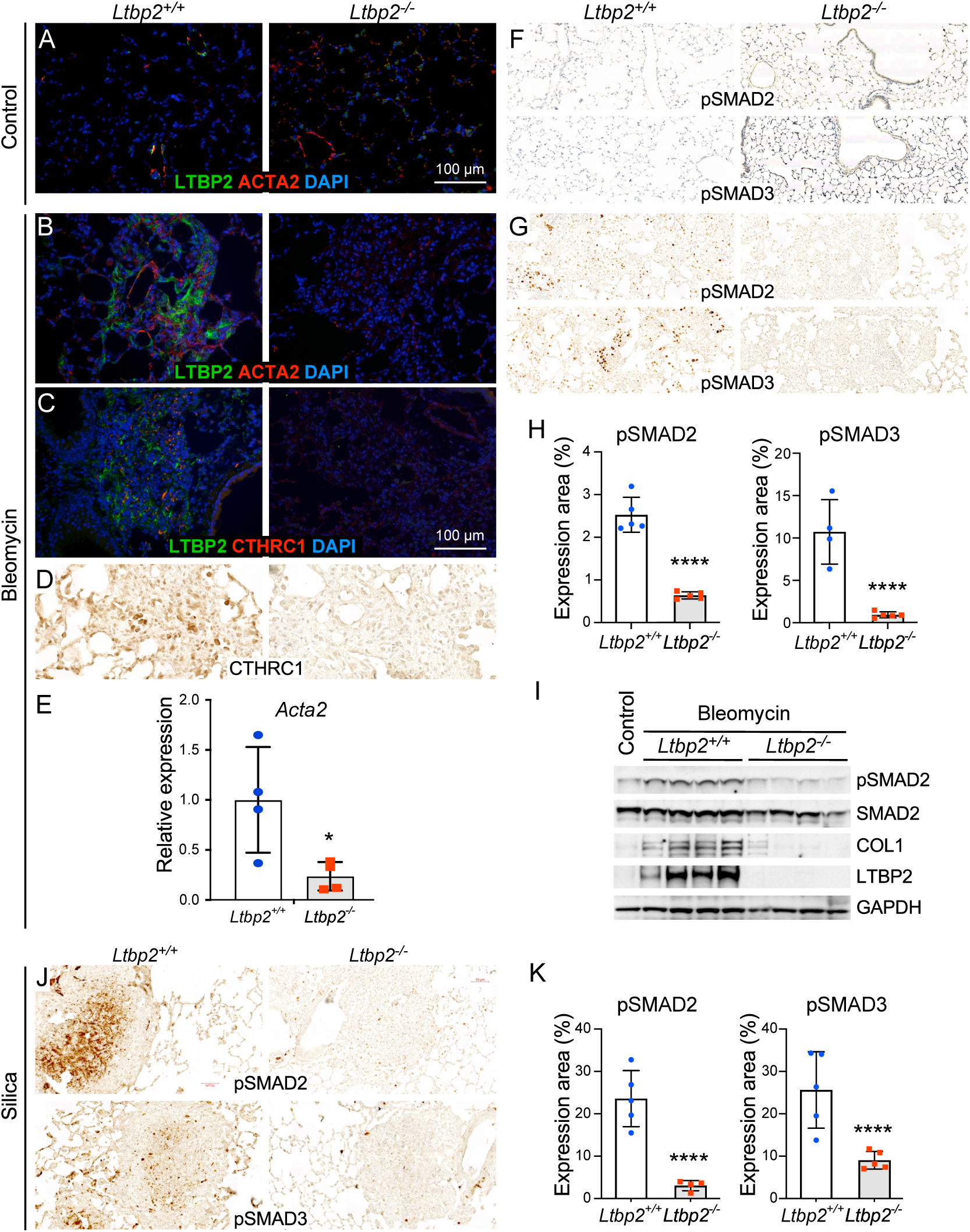
LTBP2-null mice exhibit reduced TGFB signaling in fibrosis models. (A) Immunofluorescence of lungs from untreated mice with LTBP2 (green), ACTA2 (red), and DAPI (blue) compared in WT (L panel) and *Ltbp2*^-/-^ (R panel) mice. (B) Immunofluorescence of lungs from bleomycin treated mice (28 d) with LTBP2 (green), ACTA2 (red), and DAPI (blue) compared in WT (L) and *Ltbp2*^-/-^ (R) mice. (C) Immunofluorescence of lungs from bleomycin-treated mice (28 d) with LTBP2 (green), CTHRC1 (red), and DAPI (blue) compared in WT (L) and *Ltbp2*^-/-^ (R) mice. (D) Representative CTHRC1 immunohistochemistry in bleomycin-treated *Ltbp2*^-/-^ (L panel) and control (R panel) mice. (E) Bar graph with relative expression by qRT-PCR of *Acta2* and *Pdgfra* transcripts in WT vs. *Ltbp2*^-/-^ mouse lungs at 28 d post-bleomycin. (F) Representative pSMAD2 and pSMAD3 immunohistochemistry in lungs from untreated WT (L panel) and *Ltbp2*^-/-^ (R panel) mice. (G) Representative pSMAD2 and pSMAD3 immunohistochemistry in lungs from bleomycin-treated WT (L panel) and *Ltbp2*^-/-^ (R panel) mice at 28 d. (H) Bar graph with quantification of pSMAD2 or 3 expression as percent total slide area for each mouse (n=5 per group). (I) Western blot showing expression of pSMAD2, total SMAD2, collagen I, LTBP2, and GAPDH in an untreated WT mouse (L lane) and in bleomycin-treated lungs at 28 d from WT and *Ltbp2*^-/-^ mice. (J) Representative pSMAD2 (upper panels) and pSMAD3 (lower panels) immunohistochemistry in lungs from silica-treated WT (left panels) and *Ltbp2*^-/-^ (right panels) mice at 28 d. (K) Bar graph with quantification of pSMAD2 or 3 expression as percent nodule area for each mouse (n=5 per group).

### LTBP2 enhances migration of isolated fibroblasts

Observing that LTBP2 *in vivo* serves to enhance TGFβ signaling, we next sought to evaluate its effects on TGFβ-dependent phenotypes in cultured fibroblasts. Using a scratch wound-closure assay to evaluate cell migration, a significant reduction in closure rate in primary fibroblasts isolated from *Ltbp2^-/-^* mice at 21 d post-bleomycin was seen compared with wild type (WT) controls (Fig 4A, B). Addition of a pharmacologic TGFβ receptor inhibitor further reduced migration of fibroblasts from *Ltbp2*^-/-^ mice at 48 h, suggesting residual TGFβ activity in the LTBP2 null state, while exogenous active TGFβ partially rescued the migration phenotype in *Ltbp2*^-/-^ mice (Fig 4C). LTBP2-targeted antibody reduced the rate of fibroblast migration with effect size similar to genetic deletion (Fig 4D).

**Figure 4.**
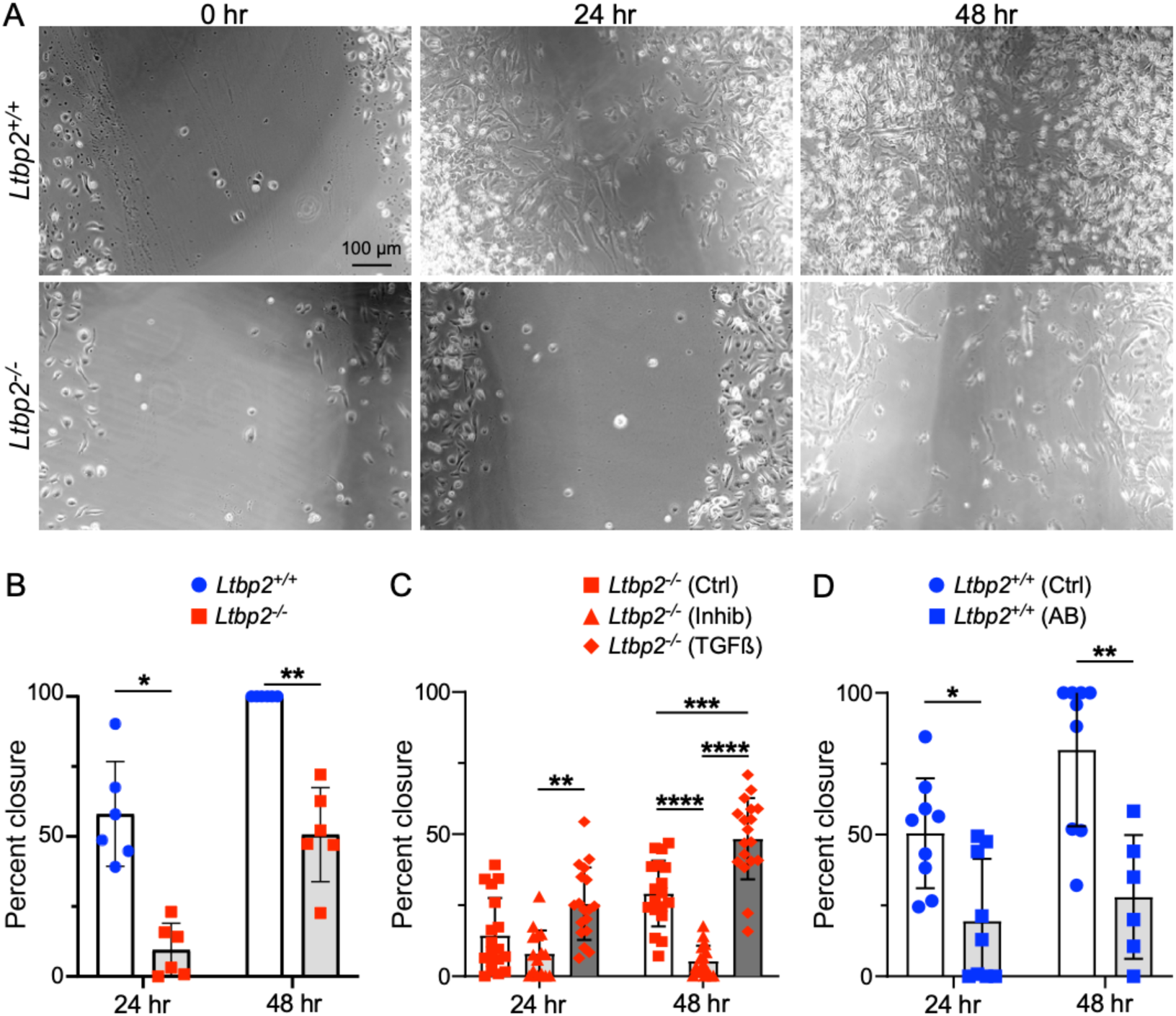
Migration is impaired in LTBP2-/- fibroblasts. (A) Representative images of scratch wound closure for WT and *Ltbp2*^-/-^ primary lung fibroblasts. (B) Bar graph depicting percent closure of scratch wound at 24 and 48 h, each point represents averaged percent closure per in three wells of primary fibroblasts from one mouse, WT versus *Ltbp2*^-/-^. (C) Bar graph depicting comparing closure of scratch wound at 24 and 48 h for *Ltbp2*^-/-^ primary fibroblasts without treatment, with TGFβ inhibitor, or with exogenous TGFβ. (D) Bar graph of scratch wound closure at 24 and 48 h for WT fibroblasts with and without treatment with anti-LTBP2 antibody.

### RNASeq of LTBP2-null and WT mice treated with bleomycin suggests loss of airway cells and increased expression of *Airn*

To identify LTBP2-dependent pathways in fibrosis, bulk RNASeq was carried out on fibrotic lungs from WT and *Ltbp2*^-/-^ mice. The 21 d timepoint was selected to minimize the contribution of fibrosis resolution to gene expression changes. Genes significantly underrepresented in *Ltbp2*^-/-^ mice after bleomycin injury included multiple markers of airway epithelium including *Scgb1a1*, *Cyp2f2*, and *Dnah6.* Genes increased in *Ltbp2*^-/-^ lungs included the lncRNA *Airn*, as well as a surprising set of genes associated with fibroblasts including *Fn1*, *Fstl1*, and *Col1a1* (Fig 5A). We further validated the increase in *Airn* in *Ltbp2*^-/-^ mice by qRT-PCR (Fig 5B). We next obtained a list of airway epithelial cell marker genes from publicly available single cell RNASeq (GSE145998) mouse lung data^23^ and evaluated their scaled expression across samples. This approach confirmed a generalized decrease in airway epithelial cell markers in *Ltbp2*^-/-^ mice after bleomycin injury (Fig 5C). By the same approach we also identified an increase in fibroblast markers in the *Ltbp2*^-/-^ mice (Fig 5D), potentially indicating that fibroblast proliferation is uncoupled from fibrosis in the absence of LTBP2, though a relative increase in fibroblast genes due to loss of airway cells is also possible. Gene ontology enrichment analysis was performed for both up and downregulated genes (Log2FC > 0.5 or < −0.5). Terms associated with increased gene expression in *Ltbp2*^-/-^ mice included ECM organization and IGFR signaling regulation (Fig 5E). Genes decreased in *Ltbp2*^-/-^ mice were associated with ciliated cells and mucosal immunity (Fig 5F). This difference was not observed in untreated *Ltbp2*^-/-^ vs. control mice (Fig S1C).

**Figure 5.**
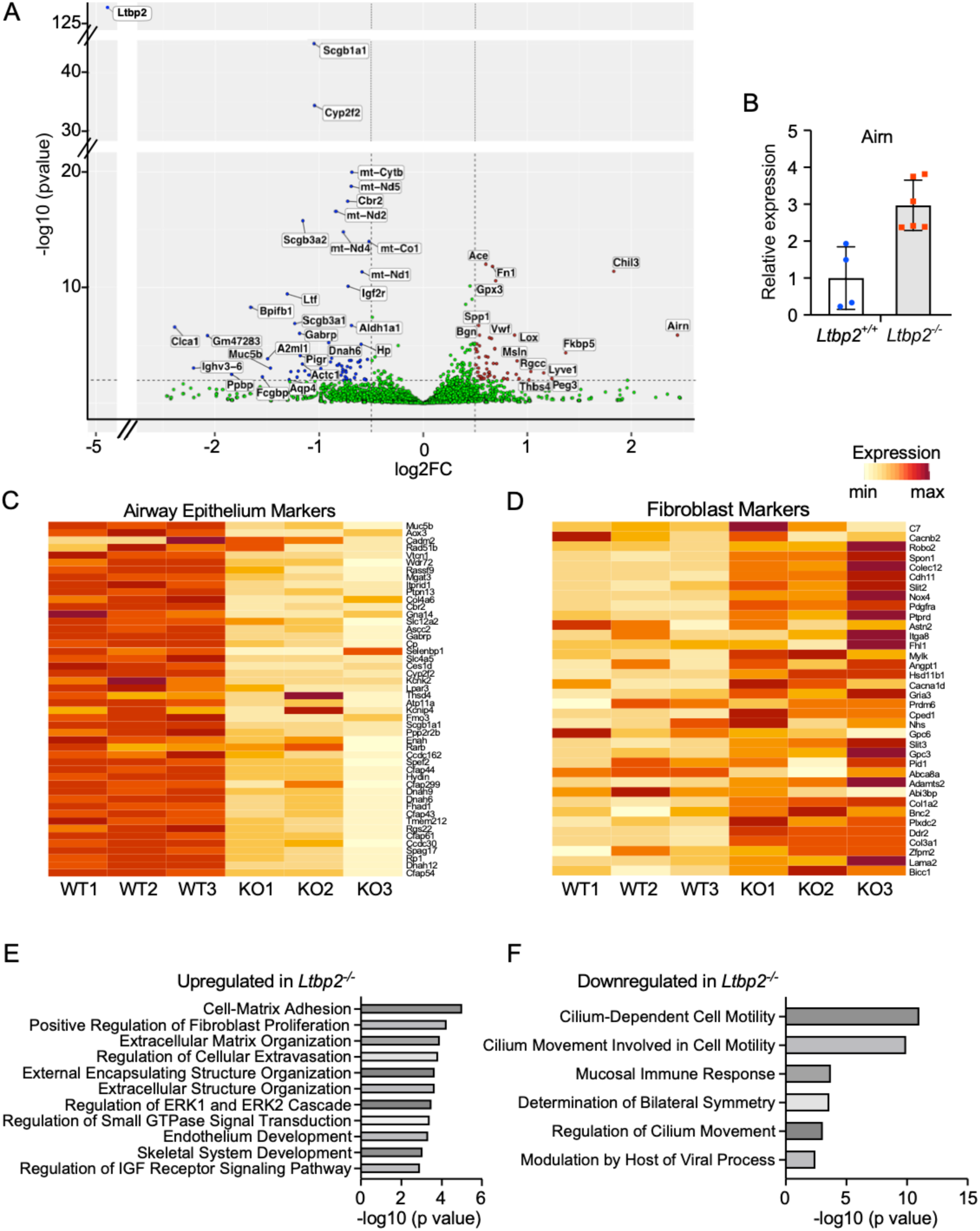
mRNA for airway genes are decreased in LTBP2 null mice after bleomycin. (A) Volcano plot of genes up- and down-regulated in *Ltbp2^-/-^* mice after 21 d bleomycin. (B) Validation of Airn upregulation in a separate cohort of *Ltbp2^-/-^* and WT mice by qRT-PCR. (C) Heatmap of top airway cell type markers (club, goblet, ciliated) derived from scRNASeq data; genes individually scaled. (D) Heatmap with expression in WT vs *Ltbp2^-/-^* mice of top fibroblast cell type markers (adventitial and alveolar fibroblasts) derived from scRNASeq data; genes individually scaled. (E) Gene ontology of top genes upregulated in *Ltbp2^-/-^* mice (logFC > 0.5) after 21 d bleomycin. (F) Gene ontology of top genes downregulated in *Ltbp2^-/-^* mice (logFC < −0.5) after 21 d bleomycin.

### *Ltbp2*^-/-^ mice exhibit deficiencies in airway epithelial repair

Airway injury has been documented previously in the bleomycin model^24^ and notably, secretory cell proportion is known to be increased in IPF^25,26^. Given the decrease in airway marker expression in our bulk RNASeq data, we performed additional immunostaining for airway markers in untreated and bleomycin-injured lungs at days 10 and 28. In WT mice, ciliated and club cell markers are abundant in airways in both conditions, whereas in *Ltbp2*^-/-^ mice there is reduction in acetylated tubulin staining after bleomycin injury (Fig 6A-C), suggesting impaired differentiation of airway progenitors to ciliated cells in the absence of LTBP2. We did not see a change in club cell number at day 28 in *Ltbp2*^-/-^ mice despite the prominence of club cell markers in our RNASeq data (Fig 6C); this may reflect the difference in selected time point or indicate a phenotypic change in these cells rather than a reduction in number. As in lung parenchyma, this airway phenotype coincided with a reduction in pSMAD2 staining in airway at day 28 post-bleomycin (Fig 6D,E), suggesting that TGFβ signaling may drive normal epithelial repair after injury. In the silica model, which is not known to be associated with airway injury, we observed no change in airway cell composition in the fibrotic lungs of *Ltbp2*^-/-^ mice (Fig 6F). To further evaluate the role of LTBP2 in airway repair we treated mice with naphthalene to selectively injure and deplete airway secretory cells. As expected, club cell sloughing was observed histologically post-naphthalene in both groups (Fig 6G), but at post-injury time points there was a reduction in CC10/Scgb1a1+ airway cells in *Ltbp2*^-/-^ mice (Fig 6G,H), consistent with deficient epithelial repair. The response to naphthalene injury was further evaluated by staining for Ki67, which showed increased airway epithelial cell proliferation at day 3 (Fig S2).

**Figure 6.**
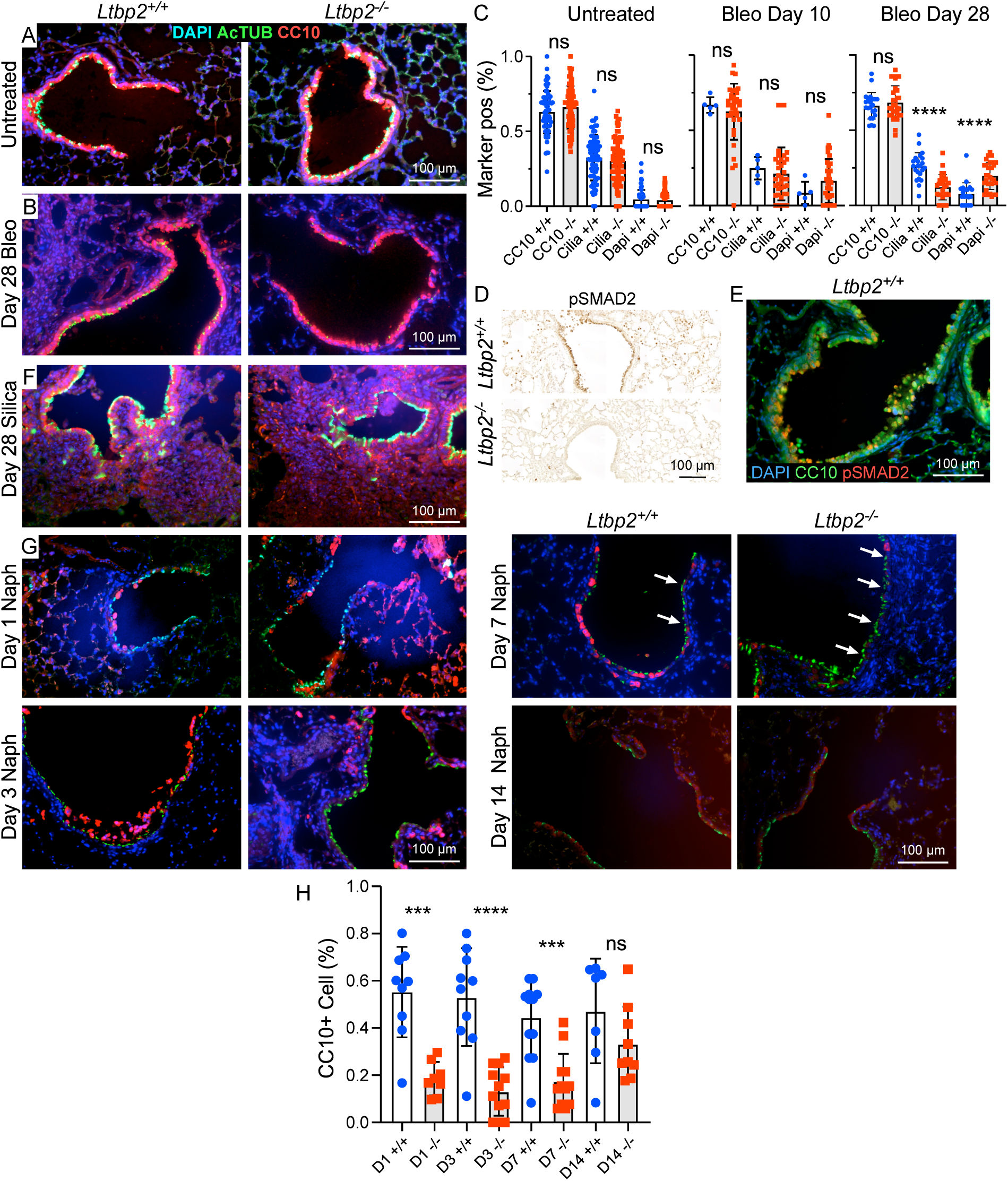
LTBP2 is required for airway epithelial repair after injury. (A) Representative immunofluorescence images of airway epithelium (club cells marked by CC10/Scgb1a1, red, and ciliated cells by acetylated tubulin, green) from untreated mice, *Ltbp2*^-/-^ and WT. (B) Representative immunofluorescence images of airway epithelium (club cells marked by CC10/Scgb1a1, red, and ciliated cells by acetylated tubulin, green) from bleomycin-treated mice at d 28, *Ltbp2*^-/-^ and WT. (C) Quantification of marker positive cells by immunostaining at 0 d (left), d10 d (middle), and 28 d (right) post bleomycin (Bleo) treatment in *Ltbp2*^-/-^ and WT mice. (D) Representative immunohistochemistry for pSMAD2 in 28 d bleomycin airways of *Ltbp2*^-/-^ and WT mice. (E) Immunofluorescence demonstrating localization of pSMAD2 (red) relative to CC10/Scgb1a1 (green) in 28 d bleomycin airway, WT only. (F) Representative immunofluorescence images of airway epithelium (club cells marked by CC10, red, and ciliated cells by acetylated tubulin, green) from untreated and silica-treated (28 d) mice, *Ltbp2*^-/-^ and WT. (G) Representative immunofluorescence images of airways after naphthalene administration in *Ltbp2*^-/-^ versus WT mice, 1, 3, 7 and 14 days, showing loss of CC10+ club cells (red) and persistence of AcTub+ ciliated cells (green). (H) Quantification of CC10+ cells in naphthalene injury model over time in *Ltbp2*^-/-^ versus WT mice.

## Discussion

Secreted matrix-binding proteins are critical regulators of growth factor signaling and their expression in the injured lung may facilitate fibrogenic responses. The microfibril-binding protein LTBP2 is of specific interest as a clinically relevant biomarker predictive of disease progression in pulmonary fibrosis, and here we employ an *Ltbp2^-/-^* mouse to evaluate the role of this protein in lung fibrosis models. We show that LTBP2 is expressed in IPF lungs and remodeled airways of COPD and asthma, that *Ltbp2* deletion is associated with decreased fibrosis and reduced TGFβ signaling, and that *Ltbp2* expression is required for effective airway repair after injury. These lung-specific features are consistent with the proposed value of LTBP2 as a biomarker for lung fibrosis.

We found that LTBP2 expression is a feature of tissue remodeling in diverse lung pathologies. These results build on previous reports showing increased protein-level expression in IPF and in small airways of COPD lungs^9,27^. We additionally observed increased LTBP2 expression in the airway walls of severe asthma, and further defined the cell types responsible for LTBP2 production at homeostasis and in disease using publicly available scRNASeq data. Activated fibroblasts expressing the pathologic fibroblast marker *CTHRC1* are enriched for expression of LTBP2, and it appears that accumulation of these cells, rather than increased expression within other fibroblast subtypes, accounts for increased LTBP2 deposition in disease.

Despite the lack of direct TGFβ binding by LTBP2 and no apparent association of baseline LTBP2 null phenotypes with TGFβ signaling^28^, we find that LTBP2 is a potent regulator of TGFβ activity in the context of lung fibrosis. Previous biochemical studies indicate that LTBP2 competes for LTBP1 (and therefore the large latent complex, LLC) binding to fibrillin-1, such that LTBP2 expression may limit TGFβ sequestration in the ECM^7^. In a fibrogenic environment the LLC may instead bind the fibronectin network via LTBP1 and undergo activation by integrins^28,29^. In addition, LTBP2 has a structural role in microfibril bundling and the formation of mature elastic fibers, the burden of which is associated with worse outcomes in IPF^30,31^. As increased matrix stiffness promotes TGFβ activation and responsiveness^32^, this may be another mechanism by which LTBP2 expression facilitates TGFβ signaling. Intriguingly, we found that treatment of cultured fibroblasts from bleomycin-treated mice with anti-LTBP2 antibody impaired migration in a similar manner to LTBP2 gene deletion, suggesting that LTBP2 targeting by antibody may have salutary effects in fibrosis. While targeting of TGFβ clinically has led to excessive inflammation and poor outcomes^33^, indirectly facilitating LLC sequestration in matrix or altering mechanical stress at sites of developing fibrosis may avoid these pitfalls.

A prior study by Zou et al. showed that LTBP2 knockdown by *in vivo* lentiviral transduction of LTBP2 shRNA was protective against bleomycin-induced fibrosis and that LTBP2 promoted myofibroblast transition in cultured fibroblasts^10^. The effect of LTBP2 was attributed to signaling via NF-kB, as decreased p65-subunit phosphorylation was seen in bleomycin-treated mice after LTBP2 knockdown. Notably, NF-kB pathway activation can occur downstream of TGFβ signaling in fibrosis models and the latter was not evaluated^34^. Our data suggests that the canonical TGFβ SMAD dependent pathway is also regulated by LTBP2. Zou et al. also observed partial inhibition of TGFβ-mediated myofibroblast transition in cultured fibroblasts by LTBP2 knockdown. This is consistent with our observed partial rescue of cell migration capacity with exogenous (active) TGFβ and suggests additional mechanisms of LTBP2-mediated TGFβ signaling beyond promoting activation of the latent form.

Increased expression of the long noncoding RNA *Airn* and concomitant downregulation of its *cis* regulatory target *Igf2r* was observed by bulk RNASeq in bleomycin-treated *Ltbp2*^-/-^ mice. *Airn* is expressed in multiple isoforms that differentially silence an array of transcripts including insulin growth factor 2 binding protein 2 (*Igf2bp2*), a profibrotic factor^35^. Interestingly, *Airn* expression has antifibrotic effects in heart and liver^35,36^. Conversely, *Igf2r* has previously been associated with fibrosis in heart and skeletal muscle^37,38^. IPF fibroblasts are reported to express higher levels of *Igf2r* mRNA than controls, though in the same study in cultured cells, Igf1r was the primary mediator of Igf2-mediated profibrotic signaling^39^. In our study, *Ltbp2*^-/-^ mice additionally had increased expression of *Igf1* and *Igfbp2*, potentially indicating broader effects of LTBP2 on insulin-like growth factor signaling. Whether *Airn* regulation is related to TGFβ pathway effects of LTBP2 cannot be determined directly from our data, and assessment of whether suppression of this transcript mediates pro-fibrotic effects of LTBP2 will be a focus of future studies.

We observed a defect in epithelial repair in *Ltbp2*^-/-^ mice in the bleomycin model and found similar effects in a naphthalene injury model targeting club cells. This was associated with impaired TGFβ signaling by pSMAD in epithelial cells and increased epithelial proliferation. Consistently, prior studies have implicated TGFβ signaling in the stimulation of airway epithelial repair and suppression of epithelial cell proliferation^40,41^. While not explored, we speculate that the development of abnormal secretory cells is a feature of IPF that may be dependent on LTBP2. Deleterious roles of TGFβ have also been described in the airway during pathological airway remodeling. In asthma (a condition where we observe increased LTBP2 expression), TGFβ has been linked to goblet cell hyperplasia and increased mucosal secretions, subepithelial fibroblast proliferation and fibrosis, smooth muscle cell hyperplasia, and epithelial cell apoptosis^42^. Whether LTBP2 inhibition would be beneficial in chronic airway remodeling processes remains to be determined.

There are several limitations to our work that will require further study. First, while TGFβ signaling is reduced in *Ltbp2*^-/-^ mice we have not shown that LTBP2 promotes fibrosis specifically via TGFβ signaling, nor that the observed effects on SMAD phosphorylation are related to altered latent TGFβ distribution in matrix. The former could be studied by examining the effect of LTBP2 knockdown in TGFBR-null mice, and the latter by demonstrating re-distribution of latent TGFβ (e.g. increased association of LTBP1 with fibronectin vs. fibrillin). Our bulk RNASeq study led us to identify LTBP2 impacts on airway epithelial differentiation. How loss of LTBP2 interrupts this process was not determined, and in preliminary studies we have not observed differences in protein-level KRT5 expression, while epithelial cell proliferation appears increased in *Ltbp2*^-/-^ mice after injury. Lastly, while lungs are histologically normal in *Ltbp2*^-/-^ mice, the use of a germ line knockout does not allow us to rule out developmental effects or alterations in immune cell populations which have been observed with other microfibril-interacting proteins^43^. This possibility would be consistent with the reductions in immune cell genes that we observed in naïve *Ltbp2*^-/-^ mice, although these markers were not differentially expressed after bleomycin injury.

In conclusion, we provide evidence of an *in vivo* profibrotic role of LTBP2, mediated in part by TGFβ signaling, for an emerging biomarker of pulmonary fibrosis progression. These data suggest that LTBP2 could be a therapeutic target in tissue remodeling in the lung, though with risk of off-target effects regarding epithelial repair.

## Acknowledgements

This work was supported by ALA DA-1277949 (JRK), NIH K08 HL159418 (JRK), NIH R21 AI167415 (DMO), NIH R01 HL176901 (DMO), NIH R01 HL167732 (MC), NIH R01 HL151685 (SLB), and the Barnes Jewish Hospital Foundation (SLB).

## Notes

### Competing Interest Statement

The authors have declared no competing interest.

### Summary of Updates

New supplemental data added (Figure S2); author list updated, Figure 3 revised. Methods/supplemental methods updated.

## References

1 Nishioka, Y., Araya, J., Tanaka, Y. & Kumanogoh, A. Pathological mechanisms and novel drug targets in fibrotic interstitial lung disease. Inflammation and Regeneration 44, 34 (2024). 10.1186/s41232-024-00345-2

2 Wilgus, T. A. Growth Factor-Extracellular Matrix Interactions Regulate Wound Repair. Adv Wound Care (New Rochelle) 1, 249–254 (2012). 10.1089/wound.2011.0344

3 Taipale, J. & Keski-Oja, J. Growth factors in the extracellular matrix. Faseb j 11, 51–59 (1997). 10.1096/fasebj.11.1.9034166

4 Zhu, J. & Clark, R. A. F. Fibronectin at Select Sites Binds Multiple Growth Factors and Enhances their Activity: Expansion of the Collaborative ECM-GF Paradigm. Journal of Investigative Dermatology 134, 895–901 (2014). 10.1038/jid.2013.484

5 Herrera, J., Henke, C. A. & Bitterman, P. B. Extracellular matrix as a driver of progressive fibrosis. J Clin Invest 128, 45–53 (2018). 10.1172/jci93557

6 Vehvilainen, P., Hyytiainen, M. & Keski-Oja, J. Matrix association of latent TGF-beta binding protein-2 (LTBP-2) is dependent on fibrillin-1. Journal of cellular physiology 221, 586–593 (2009). 10.1002/jcp.21888

7 Hirani, R., Hanssen, E. & Gibson, M. A. LTBP-2 specifically interacts with the amino-terminal region of fibrillin-1 and competes with LTBP-1 for binding to this microfibrillar protein. Matrix Biol 26, 213–223 (2007). 10.1016/j.matbio.2006.12.006

8 Robertson, I. B. et al. Latent TGF-β-binding proteins. Matrix Biol 47, 44–53 (2015). 10.1016/j.matbio.2015.05.005

9 Enomoto, Y. et al. LTBP2 is secreted from lung myofibroblasts and is a potential biomarker for idiopathic pulmonary fibrosis. Clin Sci (Lond) 132, 1565–1580 (2018). 10.1042/cs20180435

10 Zou, M. et al. Latent Transforming Growth Factor-β Binding Protein-2 Regulates Lung Fibroblast-to-Myofibroblast Differentiation in Pulmonary Fibrosis via NF-κB Signaling. Front Pharmacol 12, 788714 (2021). 10.3389/fphar.2021.788714

11 Oldham, J. M. et al. Proteomic Biomarkers of Survival in Idiopathic Pulmonary Fibrosis. American Journal of Respiratory and Critical Care Medicine 209, 1111–1120 (2023). 10.1164/rccm.202301-0117OC

12 Enomoto, Y. et al. LTBP2 is secreted from lung myofibroblasts and is a potential biomarker for idiopathic pulmonary fibrosis. Clinical Science 132, 1565–1580 (2018).

13 Zou, M. et al. Plasma LTBP2 as a potential biomarker in differential diagnosis of connective tissue disease-associated interstitial lung disease and idiopathic pulmonary fibrosis: a pilot study. Clin Exp Med 23, 4809–4816 (2023). 10.1007/s10238-023-01214-x

14 Bodmer, N. K. et al. Multi-organ phenotypes in mice lacking latent TGFβ binding protein 2 (LTBP2). Developmental Dynamics (2023).

15 Choudhury, M. et al. Targeting Pulmonary Fibrosis by SLC1A5-Dependent Glutamine Transport Blockade. American Journal of Respiratory Cell and Molecular Biology 69, 441–455 (2023). 10.1165/rcmb.2022-0339OC

16 Farooq, H. et al. Molecular imaging in experimental pulmonary fibrosis reveals that nintedanib unexpectedly modulates CCR2 immune cell infiltration. EBioMedicine 110, 105431 (2024). 10.1016/j.ebiom.2024.105431

17 Guzy, R. D. et al. Pulmonary fibrosis requires cell-autonomous mesenchymal fibroblast growth factor (FGF) signaling. The Journal of biological chemistry 292, 10364–10378 (2017). 10.1074/jbc.M117.791764

18 Inoue, T. et al. Latent TGF-β binding protein-2 is essential for the development of ciliary zonule microfibrils. Hum Mol Genet 23, 5672–5682 (2014). 10.1093/hmg/ddu283

19 Hübner, R.-H. et al. Standardized Quantification of Pulmonary Fibrosis in Histological Samples. BioTechniques 44, 507–517 (2008). 10.2144/000112729

20 Matsiukevich, D. et al. Characterization of a robust mouse model of heart failure with preserved ejection fraction. Am J Physiol Heart Circ Physiol 325, H203–h231 (2023). 10.1152/ajpheart.00038.2023

21 Tsukui, T. et al. Collagen-producing lung cell atlas identifies multiple subsets with distinct localization and relevance to fibrosis. Nature Communications 11, 1920 (2020). 10.1038/s41467-020-15647-5

22 Watanabe, N. et al. Anomalous Epithelial Variations and Ectopic Inflammatory Response in Chronic Obstructive Pulmonary Disease. Am J Respir Cell Mol Biol 67, 708–719 (2022). 10.1165/rcmb.2021-0555OC

23 Koenitzer, J. R., Wu, H., Atkinson, J. J., Brody, S. L. & Humphreys, B. D. Single-Nucleus RNA-Sequencing Profiling of Mouse Lung. Reduced Dissociation Bias and Improved Rare Cell-Type Detection Compared with Single-Cell RNA Sequencing. American Journal of Respiratory Cell and Molecular Biology 63, 739–747 (2020). 10.1165/rcmb.2020-0095MA

24 Guzy, R. D., Stoilov, I., Elton, T. J., Mecham, R. P. & Ornitz, D. M. Fibroblast growth factor 2 is required for epithelial recovery, but not for pulmonary fibrosis, in response to bleomycin. Am J Respir Cell Mol Biol 52, 116–128 (2015). 10.1165/rcmb.2014-0184OC

25 Stancil, I. T. et al. Pulmonary fibrosis distal airway epithelia are dynamically and structurally dysfunctional. Nat Commun 12, 4566 (2021). 10.1038/s41467-021-24853-8

26 Seibold, M. A. et al. The idiopathic pulmonary fibrosis honeycomb cyst contains a mucocilary pseudostratified epithelium. PLoS One 8, e58658 (2013). 10.1371/journal.pone.0058658

27 Brandsma, C. A. et al. A large lung gene expression study identifying fibulin-5 as a novel player in tissue repair in COPD. Thorax 70, 21–32 (2015). 10.1136/thoraxjnl-2014-205091

28 Rifkin, D. et al. The role of LTBPs in TGF beta signaling. Developmental Dynamics 251, 75–84 (2022). 10.1002/dvdy.331

29 Dallas, S. L. et al. Fibronectin Regulates Latent Transforming Growth Factor-beta (TGF-beta) by Controlling Matrix Assembly of Latent TGF beta-binding Protein-1. Journal of Biological Chemistry 280, 18871–18880 (2005). 10.1074/jbc.M410762200

30 Enomoto, N. et al. Amount of elastic fibers predicts prognosis of idiopathic pulmonary fibrosis. Respiratory Medicine 107, 1608–1616 (2013). 10.1016/j.rmed.2013.08.008

31 Hirai, M. et al. Latent TGF-beta-binding protein 2 binds to DANCE/fibulin-5 and regulates elastic fiber assembly. Embo j 26, 3283–3295 (2007). 10.1038/sj.emboj.7601768

32 Verma, B. K., Chatterjee, A., Kondaiah, P. & Gundiah, N. Substrate Stiffness Modulates TGF-β Activation and ECM-Associated Gene Expression in Fibroblasts. Bioengineering (Basel) 10 (2023). 10.3390/bioengineering10090998

33 Raghu, G. et al. A Phase IIb Randomized Clinical Study of an Anti-α(v)β(6) Monoclonal Antibody in Idiopathic Pulmonary Fibrosis. Am J Respir Crit Care Med 206, 1128–1139 (2022). 10.1164/rccm.202112-2824OC

34 Liu, B. et al. Costunolide inhibits pulmonary fibrosis via regulating NF-kB and TGF-β1/Smad2/Nrf2-NOX4 signaling pathways. Biochemical and Biophysical Research Communications 510, 329–333 (2019). 10.1016/j.bbrc.2019.01.104

35 Peng, T. et al. LncRNA Airn alleviates diabetic cardiac fibrosis by inhibiting activation of cardiac fibroblasts via a m6A-IMP2-p53 axis. Biology Direct 17, 32 (2022). 10.1186/s13062-022-00346-6

36 Chen, T. et al. LncRNA Airn maintains LSEC differentiation to alleviate liver fibrosis via the KLF2-eNOS-sGC pathway. BMC Med 20, 335 (2022). 10.1186/s12916-022-02523-w

37 Wu, Z. et al. Myocardial IGF2R is a critical mediator of inflammation and fibrosis after ischemia-reperfusion injury. bioRxiv (2023). 10.1101/2023.04.21.537835

38 Bella, P. et al. Blockade of IGF2R improves muscle regeneration and ameliorates Duchenne muscular dystrophy. EMBO Mol Med 12, e11019 (2020). 10.15252/emmm.201911019

39 Garrett, S. M., Hsu, E., Thomas, J. M., Pilewski, J. M. & Feghali-Bostwick, C. Insulin-like growth factor (IGF)-II- mediated fibrosis in pathogenic lung conditions. PLoS One 14, e0225422 (2019). 10.1371/journal.pone.0225422

40 Lechapt-Zalcman, E. et al. Transforming growth factor-β1 increases airway wound repair via MMP-2 upregulation: a new pathway for epithelial wound repair? American Journal of Physiology-Lung Cellular and Molecular Physiology 290, L1277–L1282 (2006). 10.1152/ajplung.00149.2005

41 Semlali, A. et al. TGF-beta suppresses EGF-induced MAPK signaling and proliferation in asthmatic epithelial cells. Am J Respir Cell Mol Biol 38, 202–208 (2008). 10.1165/rcmb.2007-0031OC

42 Ojiaku, C. A., Yoo, E. J. & Panettieri, R. A., Jr. Transforming Growth Factor β1 Function in Airway Remodeling and Hyperresponsiveness. The Missing Link? Am J Respir Cell Mol Biol 56, 432–442 (2017). 10.1165/rcmb.2016-0307TR

43 Combs, M. D. et al. Microfibril-associated Glycoprotein 2 (MAGP2) Loss of Function Has Pleiotropic Effects in Vivo*. Journal of Biological Chemistry 288, 28869–28880 (2013). 10.1074/jbc.M113.497727

