## Supplemental Figures for "Latent transforming growth factor binding protein-2 (LTBP2), an IPF biomarker of clinical decline, promotes TGF-beta signaling and lung fibrosis in mice"

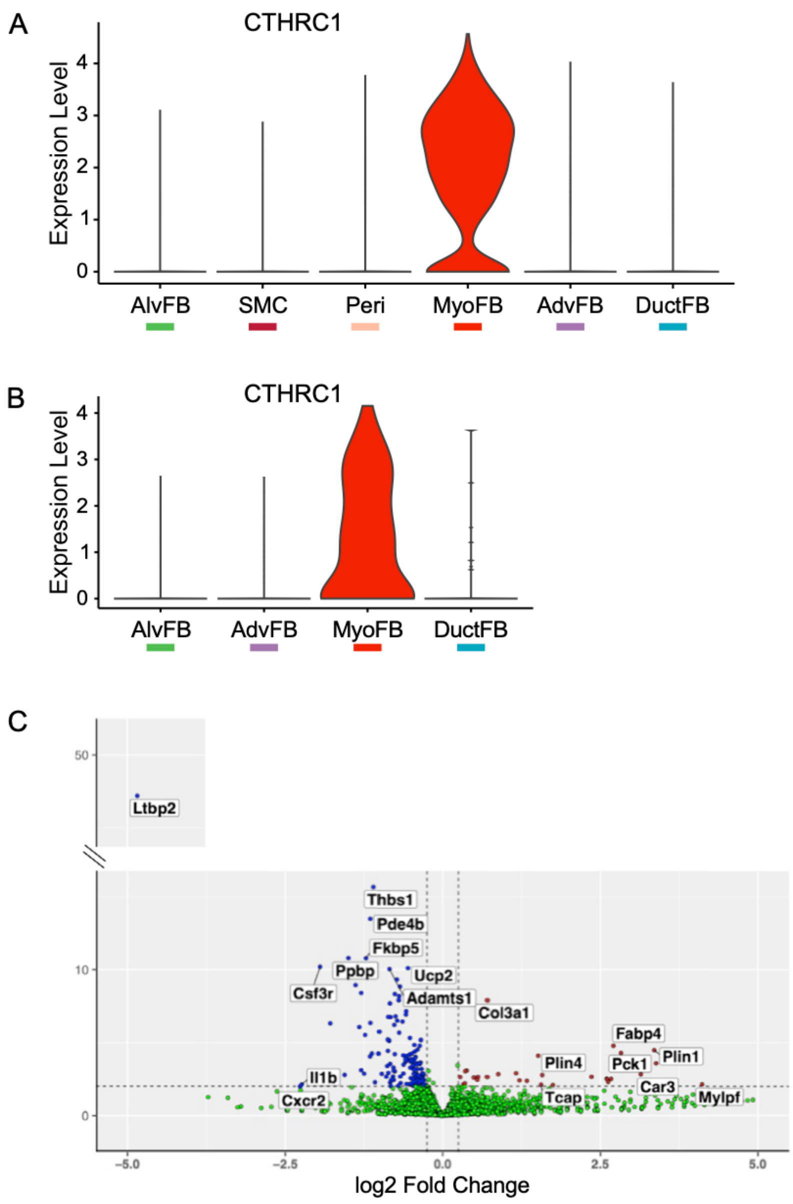


**Figure S1. Cthrc1 expression in myofibroblasts (MyoFB) from human disease scRNASeq data and differentially expressed genes from untreated Ltbp2 and WT mice by bulk RNASeq.**

A. Violin plot of *CTHRC1* expression in mesenchymal subsets from a human pulmonary fibrosis plus control dataset, Tsukui et al.

B. Violin plot of *CTHRC1* expression in fibroblast subsets from a human COPD plus control dataset.

C. Volcano plot of genes up and down-regulated in untreated *Ltbp2*$^{-/-}$ versus control mouse lungs by bulk RNASeq.

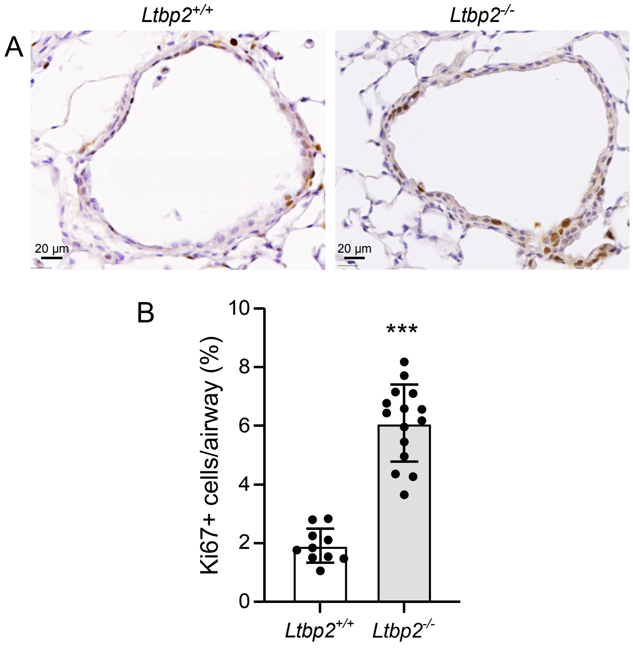


**Figure S2. Airway Ki67 expression is increased in LTBP2-/- mice after naphthalene**

A. Immunohistochemical staining of airways from *Ltbp2*$^{+/+}$ and *Ltbp2*$^{-/-}$ mice after naphthalene treatment, d3.

B. Bar plot quantifying manually counted Ki67+ cells as percentage of hematoxylin+ nuclei in each airway.
