## Supplemental Methods for "Latent transforming growth factor binding protein-2 (LTBP2), an IPF biomarker of clinical decline, promotes TGF-beta signaling and lung fibrosis in mice"

*Human lung tissues*. Studies of tissues from lungs of deceased individual donated for research and those explanted during lung transplantation were approved by the Institutional Review Board at Washington University. Tissues were anonymized. Non disease donors were those unsuitable for lung transplantation. Samples were fixed in buffered formalin and embedded in paraffin. Human lung fibroblasts isolated from non-diseased lungs donated for research and from lungs explanted from patients with idiopathic pulmonary fibrosis were obtained and characterized as previously described15 and grown in culture medium containing DMEM supplemented with 10% fetal bovine serum (FBS, Hyclone Laboratories) with assays performed at passage 3-5. Cell lines were maintained at 37º C with 5% CO2.

*Mouse models of lung injury*. Studies were approved by the Institutional Animal Care and Use Committee at Washington University. Male and female C57Bl/6J wild-type mice (Jackson Labs), 8-12 weeks of age, or Ltbp2$^{-/-}$ mice (C57Bl/6 background), 20-25 grams, were used for all models. For bleomycin administration mice were sedated with intraperitoneal ketamine (120 mg/kg) and xylazine (20 mg/kg). Bleomycin (Teva, #0703-3154-01) 1.6 U/kg intranasal (15 µl in each nostril) or saline vehicle were administered. For silica-induced fibrosis, crystalline silica particles (MIN-U-SIL 5, U.S. Silica, Mill Creek, OK, USA) were heat-treated (200º C for 2 h) to minimize endotoxin, then suspended in PBS at 200 mg/ml and sonicated for 10 min. Mice were anesthetized with isoflurane and 50 µl of suspension by oropharyngeal aspiration. For targeted airway injury, mice were administered naphthalene (250 mg/kg) or corn oil vehicle by intraperitoneal injection. Naphthalene was prepared freshly in corn oil on the day of use. At variable model endpoints (bleomycin: 7, 14, 21, or 28 d, silica: 28 d, naphthalene: 1, 3, 7, or 14 d) mice were euthanized with an overdose of ketamine and xylazine, lungs were collected for protein or RNA analysis or fixed via 10% phosphate-buffered formalin to a pressure of 20 cm $H_2O$. After overnight fixation samples were dehydrated, submitted for paraffin embedding, and cut in to 5-µm sections. Samples were stained with H&E or Masson's trichrome.

*Immunohistochemistry and immunofluorescence*. Sections were deparaffinized with xylene or citric acid and rehydrated in sequential ethanol baths. Sections were prepared for immunofluorescence or colorimetric immunohistochemical analysis. Specimens for colorimetric analysis were treated with 3% $H_2O_2$ in methanol for 30 minutes to block endogenous peroxidase activity. All sections were treated overnight at 4 °C with primary anti-LTBP2 antibody provided by Tomoyuki Nakamura (Kansai Medical University, Moriguchi, Japan) or ACTA2, PDGFRA, CTHRC1, or pSMAD2/3. Sections were incubated with appropriate secondary antibody for immunofluorescence or colorimetric analysis for 1 hour at room temperature.

*Primary fibroblast culture and scratch assay*. Briefly, lungs were dissected and treated with dispase and collagenase. Lung tissue was minced on a petri dish with sterile razors, then incubated at 37 °C in a heated shaker for 1 h. Samples were passed through sterile cell strainers and then centrifuged. The pellet was resuspended and incubated on a 75mm petri dish. Media was changed every 72 h until cells were confluent. Cells were passaged and re-plated in 48 well plates in triplicate and allowed to grow to confluence, then scratched with a sterile P200 pipet tip. Defect size was measured in 3 places at t0 and every 24 h afterwards.

*RNA isolation and qPCR*. RNA was extracted from Mouse embryonic fibroblasts using Trizol and a Qiagen RNeasy Mini kit (Qiagen, Hilden, Germany). cDNA was generated by reverse transcription using the Biorad iScript cDNA kit (Bio-Rad, Hercules, CA, USA), and mRNA determined on a StepOnePlus RT-PCR system (Thermo Fisher, 4376600). Ltbp2, Acta2, and Airn expression were assayed using Life Technologies Mm01307379_m1, Mm00725412_m1, Mm00440701_m1 Taqman primers (Life Technologies, Carlsbad, CA, USA). Calculated relative gene expression was based on average cycle (Ct) value of technical triplicates, normalized to Gapdh control, and reported as fold change. Fibrosis quantification. Lung tissue sections, including all lobes, were arranged on a single glass slide, subjected to Trichrome staining and scanned on a Zeiss Axio Scan. Z1 Slide Scanner at 20× magnification. Qupath software (version 0.4.3) was used to extract TIFF files from the whole-slide images. For bleomycin-induced fibrosis, lung sections were each divided into 2-3 similar size regions, to include the entire lobe, then scored by two blind readers using the modified Ashcroft score criteria. For silica-induced fibrosis, Qupath was used to quantify fibronodular areas. Each lobe was annotated manually to exclude the area within bronchi and the percentage of lung covered by the fibrotic masses was generated by Qupath based on the density of the fibrotic nodules.

*Lung tissue assay for hydroxyproline*. Snap frozen left lungs were acid hydrolyzed and hydroxyproline was determined colorimetrically using reagents according to the manufacturer's protocol (Sigma–Aldrich, #MAK008).

*RNA Sequencing*. Snap frozen lung tissue was stored at −80 °C until processing. mRNA isolation was performed with the Ambion PureLink RNA Mini Kit (Thermo Fisher, 12183018, Lot 2379554), and samples submitted for library preparation and sequencing through the Washington University Genome Technology Access Center (GTAC, https://gtac.wustl.edu/). RNA quality was determined using an Agilent Bioanalyzer, then library preparation performed with 0.5–1 μg of total RNA. After removal of rRNA using a RiboErase kit (Kapa Biosystems), mRNA was fragmented in reverse transcriptase buffer and heated to 94°C for 8 min. mRNA reverse transcription was performed using SuperScript III RT enzyme (Thermo Fisher-Life Technologies, per manufacturer's instructions) and random hexamers to generate cDNA, followed by a second strand reaction to yield ds-cDNA. Illumina sequencing adapters were then appended as per manufacturer protocol, and ligated fragments amplified for 12–15 cycles using unique dual index primers. Sequencing was performed on an Illumina NovaSeq-6000 flow cell using paired-end reads extending 150 bases. For read processing, raw reads were first trimmed using Cutadapt (v.3.5) to remove low-quality bases and reads. Trimmed reads were then aligned to the mouse genome mm10 with GENCODE annotation vM25 using STAR (v.2.7.9) with default parameters. Transcript quantification was performed using featureCounts from the Subread package (v.2.0.1), with further quality control assessments made using RSeQC (v.4.0.0) and RSEM (v.1.3.1). Batch correction was performed using edgeR, EDASeq, and RUVSeq (Bioconductor version: release 3.16). Principal component analysis (PCA) and differential expression analysis for samples were determined using DESeq2 in negative binomial mode using batch-corrected transcripts from feature counts (>2-fold expression change, >1.5 counts/million, and Benjamini corrected $P < 0.05$). Pairwise comparisons were made between control and Ltbp2-/- mice to identify differentially expressed genes (DEGs). Gene ontology (GO) analyses were performed using Enrichr (https://maayanlab.cloud/Enrichr/). For selection of airway and fibroblast-specific genes, single cell RNASeq data from WT mouse lung was downloaded from GEO (GSE145998) and analyzed with *Seurat*: marker gene lists for individual cell types were obtained using *FindAllMarkers* with default parameters, and top genes for major airway epithelial subclusters (ciliated, secretory, and goblet cells) and fibroblast subclusters were used.

*Single cell RNASeq analysis*. Datasets GSE132771 (fibrosis) and GSE151974 (COPD) were accessed via the Gene Expression Omnibus (GEO). Filtered count matrices were loaded into Seurat and clustering performed using the standard pipeline. After annotation using canonical markers, subsets of mesenchymal cells were re-clustered and cell type-specific markers obtained using FindAllMarkers with default parameters. Subclusters were annotated using described markers from Tsukui et al[21].

*Statistical Analysis***.** Significant differences in mean values were calculated using paired Student's *t* tests or one-way ANOVA. Mortality was analyzed using log-rank (Mantel-Cox) test and two-way ANOVA. A *P* value of less than 0.05 was considered significant.
